# Effects of chlorpyrifos on early development and anti-predator behavior of agile frogs

**DOI:** 10.1101/2021.11.03.467073

**Authors:** Zsanett Mikó, Veronika Bókony, Nikolett Ujhegyi, Edina Nemesházi, Réka Erös, Stephanie Orf, Attila Hettyey

## Abstract

The widespread application of pesticides makes it important to understand the impacts of these chemicals on wildlife populations. Chlorpyrifos is an organophosphate insecticide which can affect the development and behavior of aquatic organisms and may thereby alter predator-prey interactions. To investigate how environmentally relevant, sublethal concentrations of chlorpyrifos affect anti-predator behavior and larval development of the agile frog (*Rana dalmatina*), we exposed tadpoles to one of three concentrations (0, 0.5 and 5 μg chlorpyrifos / L) either for a brief period of three days (acute exposure) or throughout larval development (chronic exposure). We observed tadpole activity and space use in the presence or absence of chemical cues of predatory fish. We also assessed mortality, time to metamorphosis, mass at metamorphosis, brain morphology and sex ratio. We found that tadpoles chronically exposed to 5 μg/L chlorpyrifos swam distances that were longer by more than 20 % and exhibited body masses at metamorphosis that were lower by ca. 7 % than in control individuals, but the other fitness-related traits remained unaffected. The lower concentration of chlorpyrifos applied chronically, and either one of the acute chlorpyrifos treatments did not influence any measured trait. Our results demonstrate that exposure to chlorpyrifos can induce changes in locomotor activity and may result in lowered body mass of agile frog tadpoles, but only if the insecticide is present chronically at concentrations which are rarely reached in natural waterbodies. Thus, agile frog tadpoles appear to be relatively tolerant to chlorpyrifos, but may nonetheless suffer from its presence in situations of repeated high-dose application.

## Introduction

In aquatic systems, chemical cues are important sources of information about predation risk for a wide range of taxa. When predators capture, consume and digest prey, the released chemicals can provide detailed information about predation risk (Hettyey et al., 2015; Schoeppner and Relyea, 2009), which in turn can induce adaptive (and maladaptive) plastic changes in animals (Tollrian and Harvell, 1990). Aquatic pollutants can affect several aspects of animal behavior, including change in microhabitat use (Tierney et al., 2007; Yu et al., 2014), foraging behavior (Pavlov et al., 1992; Semlitsch et al., 1995), can also cause abnormal motion (Denoël et al., 2013; Levin et al., 2004), and predator avoidance (Bridges, 1999; Scholz et al., 2000).

In aquatic toxicology, the most often used vertebrate model animals are fishes (Melvin and Wilson, 2013), standard toxicity testing and authorization procedures also requires testing on this model animals (Adams and Rowland, 2003; Nikinmaa, 2014). Studying the effects of pesticides on the behavior of amphibians is similarly important, because amphibians are an especially endangered class of vertebrates (IUCN, 2021), and they are especially susceptible to chemical pollution due to their highly permeable skin and complex life cycle. Furthermore, because many amphibians use small puddles, temporary ponds and ditches as breeding sites, which often occur adjacent to agricultural fields, they are especially likely to become exposed to high concentrations of agricultural contaminants during their larval life (Bridges, 1997), which is the most sensitive life stage due to intense development and growth, and the re-organisation of entire organ systems during metamorphosis.

Chlorpyrifos (O,O-Diethyl O-3,5,6-trichloropyridin-2-yl phosphorothioate) is a broadspectrum organophosphate insecticide applied in large quantities both in agricultural and urban settings (Bernabò et al., 2011b). Its primary mode of action is the inhibition of cholinesterases (ChEs), enzymes involved in the normal function of nerve and muscles synapses. Blocking of cholinesterases can induce neurotoxic effects in insects as well as nontarget organisms (Barron and Woodburn, 1995) via the buildup of excessive acetylcholine in the synapses, causing hyperactivity and uncontrolled muscle spasms and, eventually, paralysis, respiratory failure, and death (Barron and Woodburn, 1995; De Arcaute et al., 2012). Aquatic environments are easily contaminated by chlorpyrifos through direct application against mosquitoes, as well as spray drift and runoff from adjacent agricultural areas (Bernabò et al., 2011a; Widder and Bidwell, 2008). Chlorpyrifos concentration in surface waters generally ranges from 0.02 ng/L to 10.8 μg/L, but can be as high as 96 μg/L (Arain et al., 2018; Chowdhury et al., 2012; Claver et al., 2006; Delgado-Moreno et al., 2011; Jergentz et al., 2005; Kurt-Karakus et al., 2011; Marino and Ronco, 2005; Rico et al., 2021; Xue et al., 2005).

Chlorpyrifos can be severely toxic to aquatic invertebrates and fish, and the highest measured environmental concentrations can also cause excessive mortality in amphibian tadpoles (Carriger and Rand, 2008; Giesy et al., 1999). Furthermore, an apparent connection between the presence of chlorpyrifos residues and reductions in amphibian population sizes has been reported both at local (Fellers et al., 2004) and at landscape scale (Sparling et al., 2001). Nevertheless, we still have scarce knowledge regarding the ecotoxicological effects of environmentally relevant concentrations of chlorpyrifos on amphibians (Bernabò et al., 2011b; Giesy et al., 1999; Richards and Kendall, 2003, 2002).

In this study, our aim was to investigate the effects of chlorpyrifos on larval development and antipredator behavior of the agile frog (*Rana dalmatina*). To achieve this, we exposed tadpoles to one of two ecologically relevant concentrations of chlorpyrifos throughout their larval life, and investigated its effects on mortality, speed of somatic development, body size at metamorphosis, brain morphology, and sex ratio. To determine if brief exposure to chlorpyrifos can also cause changes in the abovementioned traits, i.e. if the insecticide is only applied once and thereafter its concentration diminishes, we also applied the chemical treatments for a brief period of three days. Finally, to assess the effect of chlorpyrifos on antipredator behavior, we observed the behavior of chlorpyrifos-exposed and control tadpoles in the presence or absence of chemical cues of predatory fish. We chose this species because it occurs and breeds near or in agricultural and urbanized areas (Nemesházi et al., 2020).

## Methods

In March 2018 we collected 50 eggs from each of ten freshly laid egg-clutches of the agile frog (*Rana dalmatina*) from a pond in the Pilis, a hilly woodland in Hungary (Szárazfarkas, 47°44’ 4.12” N, 18°49’ 7.04” E) and transported them to the Julianna-major Experimental Station of the Plant Protection Institute in Budapest (47°32’ 52” N, 18°56’ 07” E). Until hatching, we kept each clutch separately in the laboratory in 3-L containers holding 1 L of reconstituted soft water (RSW: 48 mg NaHCO_3_, 30 mg CaSO_4_ 2 H_2_O, 61 mg MgSO_4_ 7 H_2_O, 2 mg KCl added to 1 L reverse-osmosis filtered, UV sterilized tap water), at 19 °C and a photoperiod mimicking outdoors light-dark cycles (starting with 12:12 h light:dark cycles in late March, which we gradually changed to 14:10 h by the end of April). When the hatchlings reached the free swimming stage, i.e. developmental stage 25 (Gosner, 1960), we started the experiment by haphazardly selecting 26 healthy-looking tadpoles from six clutches and 32 tadpoles from four clutches (284 larvae in total) and placing them into rearing containers. We reared the tadpoles at 20° C individually in 1.5-L containers filled with 1 L RSW, arranged in a randomized block design. Twice per week, we changed the rearing water and fed the tadpoles ad libitum with slightly boiled, chopped spinach. The remaining eggs and tadpoles were released at their pond of origin.

The 284 tadpoles were distributed among 10 treatment groups as follows (Fig 1). Besides the control group (no chlorpyrifos), we applied 0.5 or 5 μg/L chlorpyrifos either acutely (i.e. for 3 days around behavioral observations, see below) or chronically (i.e. during the entire larval development), and these 5 chemical treatments were combined with the presence or absence of chemical cues of predatory fish during behavioral observations (see below). In case of the control and the chronic exposure treatments we had 34 individuals per group, whereas in the acute treatments we had 20 individuals per group. We increased sample sizes in the former treatment groups to ensure the proper number of individuals to sex ratio determination. In the control treatment, we kept tadpoles in chlorpyrifos-free RSW throughout the experiment, to which we added 1 μL of 96 % ethanol as solvent control. The resulting ethanol concentration (<0.001 ‰) was much lower than those observed to harm anuran embryos or tadpoles (0.4 % and 0.13 % respectively, Fainsod and Kot-Leibovich, 2018; Peng et al., 2005; Taylor and Brundage, 2013). The other treatment groups were exposed to one of the two nominal concentrations (i.e. 0.5 or 5 μg/L) of chlorpyrifos dissolved in ethanol. Tadpoles assigned to the chronic chlorpyrifos treatments were exposed to the insecticide from developmental stage 25 to 42 (start of metamorphic climax), whereas the acute chlorpyrifos treatments lasted for three days, starting at the beginning of the first behavioral observation on day 30 and ended 96 hours later, just before the second behavioral observation. In all treatment groups, we renewed the nominal concentrations at each water change. To measure actual concentrations in rearing containers, we took samples from the rearing water of tadpoles into amber PET flasks once per week over three weeks, taking one liter per nominal concentration (0, 0.5 or 5 μg/L chlorpyrifos) on each occasion. The samples were immediately transported to an accredited analytical laboratory (SynTech Research Hungary Kft.), and were analyzed as described in Bókony et al. (2020). These analyses detected no chlorpyrifos in the control treatment (lowest quantification level: 50 ng/L), whereas the concentrations measured in samples treated with chlorpyrifos were close to the nominal concentrations (0.533, 0.498, and 0.35 μg/L in treatments with a nominal concentration of 0.5 μg/L; 4.76, 4.13, and 5.4 μg/L in treatments with a nominal concentration of 5 μg/L). The concentrations given throughout the text henceforth are nominal.

**Fig. 1.**
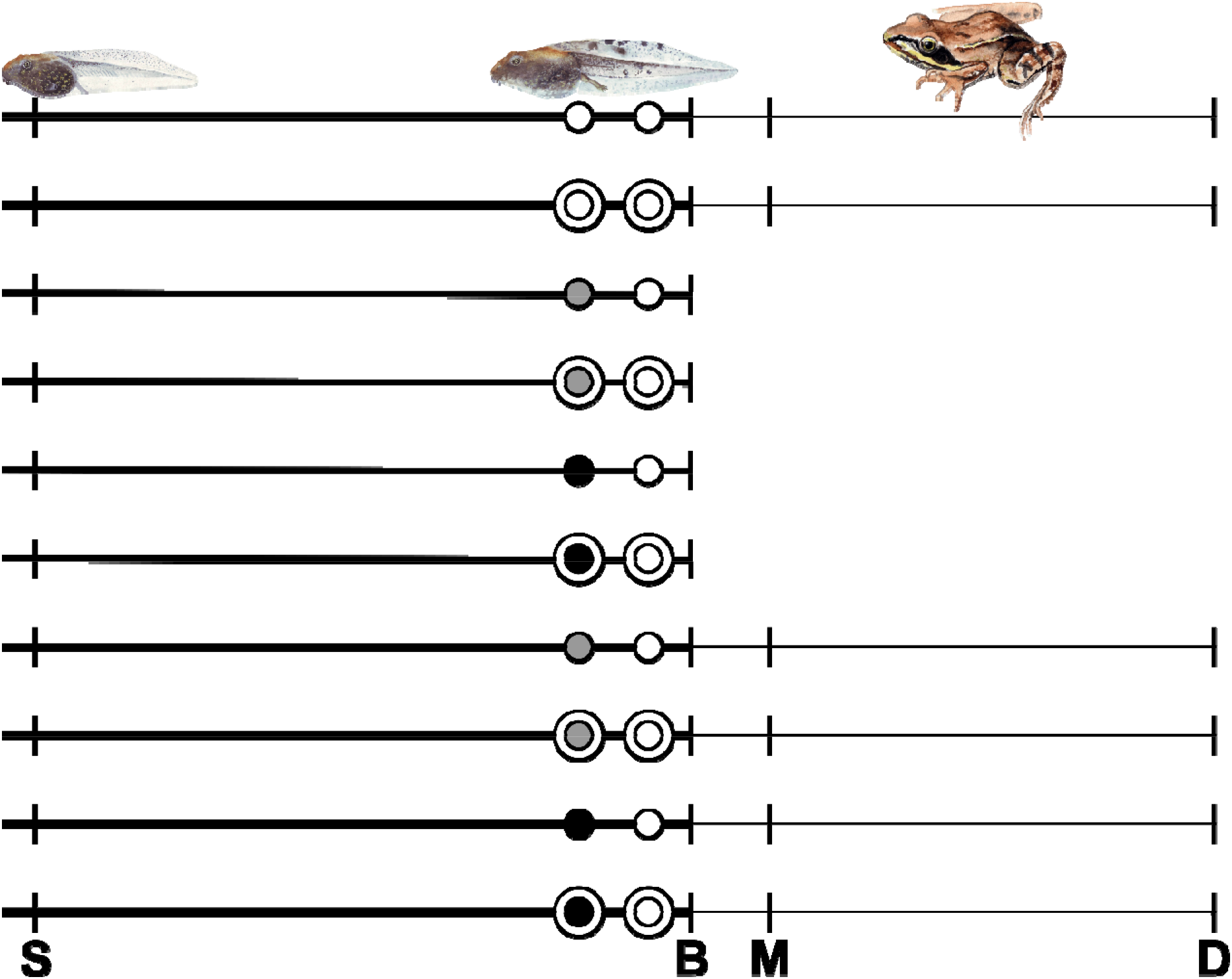
Schematic illustration of the 10 treatments, each horizontal line representing a treatment group. Longer lines are the chronic, shorter lines are the acute treatments, narrow lines represent the reduced sample size after the brain sampling. Circles symbolize behavioral observations: white circles mean that animals did not get chlorpyrifos, gray circles are the 0.5 μg/L and black circles are the 5 μg/L chlorpyrifos treatment. Double circles symbolize the predator cue. S means the start of the experiment, B is the brain sampling, M is the start of metamorphosis and D is the dissection of the animals.

Thirty days after start of the experiment we video-recorded the behavior of 200 individuals, 20 from each of the 10 treatment groups. We transferred each tadpole into a new container identical to the rearing containers but with no food, filled with 1 L RSW treated with 0, 0.5 or 5 μg/L chlorpyrifos according to their assigned treatments; tadpoles in both the acute and the chronical treatments received chlorpyrifos this time. After recording 20 minutes of baseline behavior, we added 40 ml pure RSW or 40 ml RSW containing chemical cues of predatory fish into each container, and recorded the behavior of the animals for another 20 minutes. The chemical cues were prepared as described in Hettyey et al. (2016), using water from a 130-L tank holding 4 European perch (*Perca fluviatilis*) that had been feeding on agile frog tadpoles. The fish were obtained and maintained as described in Üveges et al., (2021). As the European perch is a native predator in our region, agile frog tadpoles respond to chemical cues indicating its presence by decreasing their activity if fish had been feeding on conspecific tadpoles, even if they are predator-naïve (Hettyey et al., 2016). We used three digital camcorders (Canon LEGRIA HF R66) to record the behavior of tadpoles. Because ten rearing containers fitted under one camera, we performed seven consecutive rounds of recordings, where the order of individuals among rounds was randomized. After recording, we returned containers to the rearing shelves and we fed the tadpoles.

On day 33 we repeated the above process of recording tadpole behavior, except that this time none of the animals received chlorpyrifos during the video recording (i.e. the acute exposure ended after 3 days, right before the second observation and the chronic exposure group spent this 40 minutes without the chemical). For each individual, the predator-cue treatment was the same as during its first observation. After 40 minutes of observation, we changed the water in the chronically treated groups to chlorpyrifos-treated RSW, returned all containers to the rearing shelves, and fed the tadpoles. One day after the second behavioral observation we randomly chose 100 individuals, 10 from each treatment group, measured their body mass (± 0.1 mg), and preserved them in 4% formalin - 0.1 M phosphate-buffered saline solution for later brain measurements. Subsequently, we terminated the acute exposure treatments, and released remaining tadpoles at their pond of origin.

The remaining animals (the chronically treated groups and the controls) were raised further for measuring developmental endpoints. When a tadpole reached developmental stage 42 (start of metamorphosis, identified by the appearance of forelimbs), we replaced its rearing water with 0.1 L pure RSW (i.e. containing neither ethanol nor chlorpyrifos) and slightly tilted the container to allow the animal to leave the water. When metamorphosis was completed (developmental stage 46, identified by complete tail resorption), we moved the animal into a clean rearing box that contained wet paper towels as a substrate and a piece of egg carton as shelter, which we changed every other week. Metamorphosed animals were fed *ad libitum* with springtails and small (2-3 mm) crickets sprinkled with a 3:1 mixture of Reptiland 76280 (Trixie Heimtierbedarf GmbH & Co. KG, Tarp, Germany) and Promotor 43 (Laboratorios Calier S.A., Barcelona, Spain) containing vitamins, minerals and amino-acids.

We dissected animals 59-86 days after metamorphosis (102-131 days after they reached the free-swimming tadpole stage), when the gonads are already well differentiated in this species (Bernabò et al., 2011b; Ogielska and Kotusz, 2004). When froglets reached this age, we measured their body mass to the nearest 0.01 g and euthanized them using a water bath containing 6.6 g/L tricaine-methanesulfonate (MS-222) buffered to neutral pH with the same amount of Na_2_HPO_4_. After dissection, we cut out the entire digestive tract and measured its mass to the nearest 0.01 g, because many animals’ guts contained food although we had not fed them for 2-4 days before dissection. We examined the gonads under an Olympus SZX12 stereomicroscope at 16× magnification, and categorized phenotypic sex as male (testes) or female (ovaries) based on the gross anatomy of the gonads.

### Brain measurements

The whole brains were dissected from the cranium of the animals stored in fixative solution under a Zeiss SteREO Discovery V8 microscope, placed on glass beads, and photographed from dorsal, lateral (left) and ventral views with a Canon 600D digital camera. All photographs were analyzed using TpsDig 2.31 software (https://life.bio.sunysb.edu/ee/rohlf/software.html). For the whole brain as well as for each brain region (bulbus olfactorius, telencephalon, tectum optica, medulla oblongata, diencephalon, and hypothalamus), height, length, and width were measured as the greatest distance enclosed by the given structure in the given direction, converted to mm using a size reference included in each picture. From these three size dimensions, we estimated the volume of the brain and each brain region according to the ellipsoid model following the formulas (including the correction factor) of Pollen et al. (2007). We measured 20 randomly chosen brains three times; these measurements showed high repeatability (intra-class correlation coefficient ≥ 0.85). The medulla got damaged during dissection in two animals, so we could not estimate total brain size in these two cases.

### Video analyses

To analyze the behavior of tadpoles, we used a freely available video tracking program, ToxTrac (Rodriguez et al., 2018). We divided the observations into two parts, before and after the addition of the 40 ml stimulus water into the containers, yielding 400 recordings per day. One observer checked all recordings by comparing them with the result of the automatic tracking, and found tracking errors in 21 recordings (belongs to 13 individuals); we excluded these 21 recordings (out of 800) from the statistical analyses. We extracted the following variables from each 20-min recording: mobility rate (proportion of time spent moving), the swimming trajectory length (total distance traveled by the tadpole in kilopixels), and the proportion of time it spent close to the sides of the container (i.e. its body was within a 50-pixel wide stretch measured from the side, where the tadpoles had an average body length of ca. 40 pixels in the videos).

### Statistical analyses

We analyzed the data in the R computing environment v4.0.3 (R Core Team, 2020). For the behavioral variables we used linear mixed-effects models (‘lmer’ function of the ‘lme4’ package), where the behavioral variable after stimulus was the dependent variable, the chlorpyrifos treatment (5 groups), the predator treatment (presence/absence of chemical cues of predatory fish), and the day of observation (first or second) were the fixed factors, and individual nested within family were the random factors. We also included the behavioral variables before stimulus (i.e. in the first 20 minutes) and the time of day (observation session, ranging from 1 to 7) as numeric covariates, and all two-way interactions between the fixed factors as well as their three-way interaction.

In the analysis of total brain size, the fixed effects were the chlorpyrifos treatments (5 groups) and body mass was a covariate. These same predictors were included in the models of each brain region, where we entered total brain size instead of body mass as a covariate. We graphically examined the relationship between total brain size and body mass, and also between each brain region and total brain size, both with and without logarithmically transforming the variables, and we found no indication that power-function relationships would fit the data better than linear relationships, so we used the latter (Pollen et al., 2007).

For the analysis of survival, we used a generalized linear model with binomial distribution and bias-reduction adjustment for separation (‘glm’ function of the ‘brglm2’ package). We could not apply Cox regression in this case, because in some treatment groups all individuals survived until dissection, thus the model cannot compute survival rate. In this analysis, survival was treated as a binary variable, expressing whether the individual survived to dissection. To analyze body mass at metamorphosis and body mass at dissection we used linear mixed-effects models (‘lmer’ function of the ‘lme4’ package). For the analysis of time to metamorphosis we used generalized linear mixed model with Conway-Maxwell Poisson distribution (‘glmmTMB’ function of the ‘glmmTMB’ package), and generalized mixed-effects model with binomial distribution for sex ratio (‘glmer’ function of the ‘lme4’ package). Because individuals were exposed to predator chemical cues for only a short period, we did not enter this factor into the analyses of the fitness-related traits. Only chemical treatment (3 groups: control and chronic exposures) was tested as a fixed factor in these models, and family as random factor (except for survival, because the bias-reduced analysis cannot accommodate random effects). In case of body mass at dissection we also entered days from the start of the experiment to dissection as a covariate.

## Results

Out of the three behavioral variables, only the swimming trajectory length was affected by chlorpyrifos (Table 1): locomotor activity increased in tadpoles exposed to the high concentration chronically, while no change was detectable in any other treatment group (Fig. 2). The presence of predator chemical cues had no significant effect on behavior except for the position of the tadpoles within the containers (Table 1): individuals that received predator cues spent less time along the edges (mean ± SE: 70.7 ± 1.47 %) than individuals that received no predator cues (mean ± SE: 74.8 ± 1.46 %). The interactions between chemical treatment, predator treatment, and observation day were all non-significant for all three behavioral variables (Table 1, Table S1).

**Table 1.**
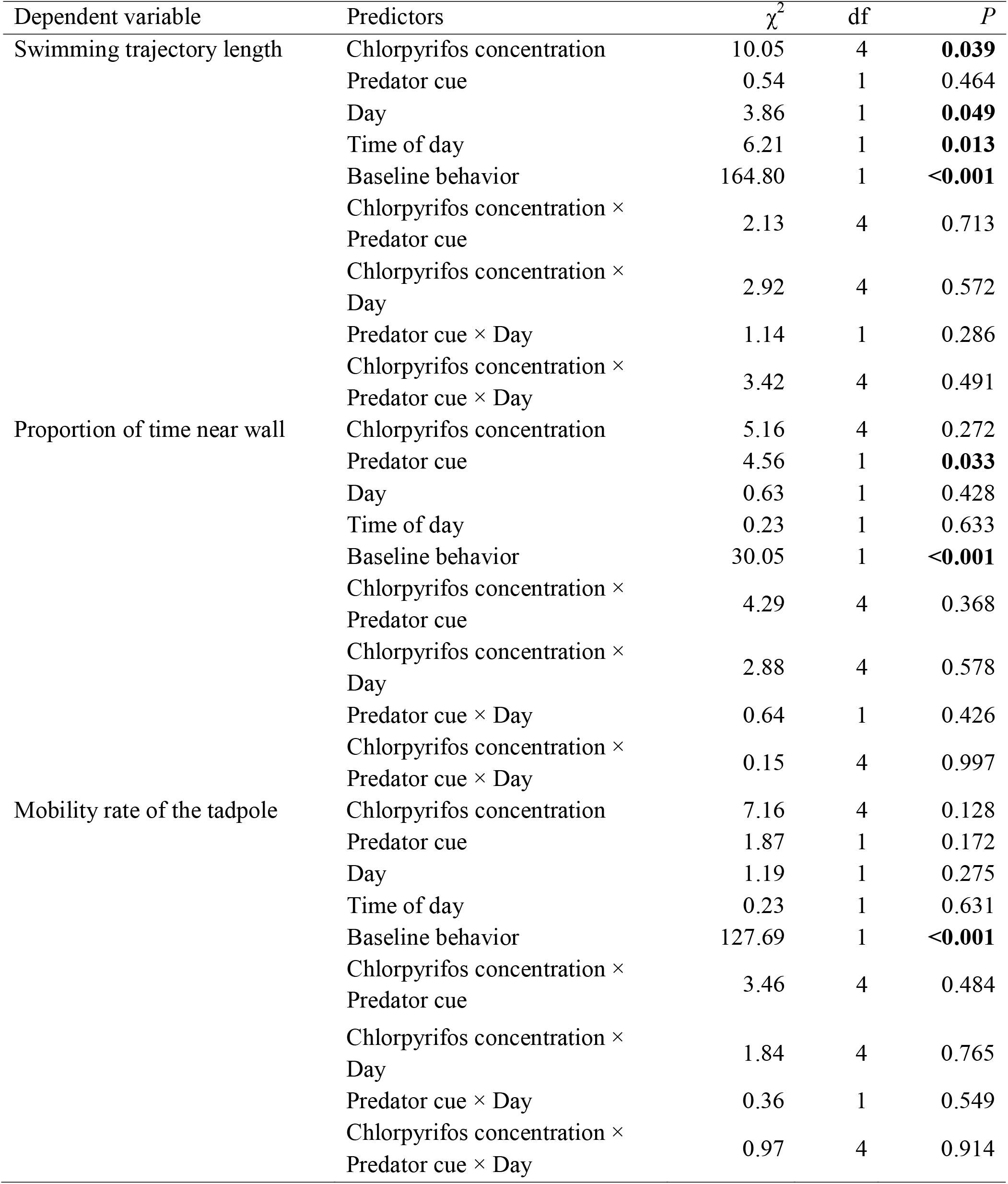
Type-2 analysis-of-deviance tables of the effects of acute and chronic exposure to chlorpyrifos at a concentration of 0.5 and 5 μg/L on the behavioral variables in the presence or asence of predator cues on the first and second observation day. Significant effects (*P* < 0.05) are highlighted in bold. Baseline behavior means the first 20 minutes of recording without predator cue. Time of day is the observation session, ranging from 1 to 7.

**Fig 2.**
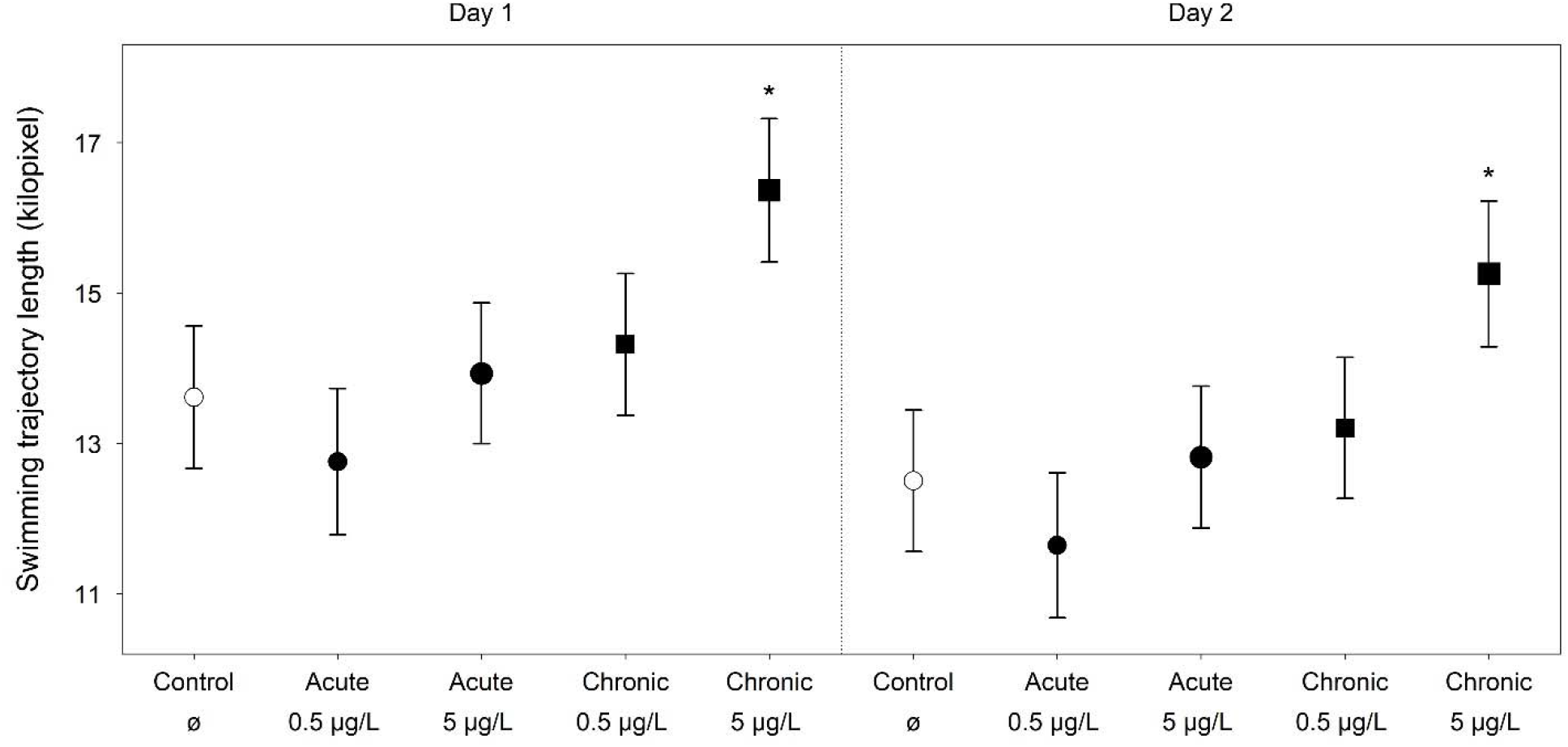
Distances traveled by tadpoles after the addition of stimulus water in each chlorpyrifos treatment group (regardless of presence or absence of predator cues). Error bars show the means and standard errors estimated from the model in Table 1.

Total brain size was positively correlated with body mass, where larger tadpoles had larger brains. However, exposure to chlorpyrifos had no effect either on total brain size or on the size of the brain regions (Table 2).

**Table 2.** Type-2 analysis-of-deviance tables of the effects of acute and chronic exposure to chlorpyrifos at a concentration of 0.5 and 5 μg/L on the size of total brain and each brain region. Significant effects (*P* < 0.05) are highlighted in bold. Body mass was measured just before sample collection.

| Dependent variable | Predictors | $\chi^2$ | df | *P* |
| --- | --- | --- | --- | --- |
| Total brain size | Chlorpyrifos concentration | 1.77 | 4 | 0.779 |
|  | Body mass | 10.83 | 1 | **0.001** |
| Bulbus olfactorius | Chlorpyrifos concentration | 5.49 | 4 | 0.239 |
|  | Total brain size | 11.05 | 1 | **<0.001** |
| Telencephalon | Chlorpyrifos concentration | 4.35 | 4 | 0.361 |
|  | Total brain size | 20.21 | 1 | **<0.001** |
| Tectum optica | Chlorpyrifos concentration | 1.24 | 4 | 0.872 |
|  | Total brain size | 41.33 | 1 | **<0.001** |
| Medulla oblongata | Chlorpyrifos concentration | 5.87 | 4 | 0.209 |
|  | Total brain size | 63.35 | 1 | **<0.001** |
| Diencephalon | Chlorpyrifos concentration | 3.09 | 4 | 0.544 |
|  | Total brain size | 36.50 | 1 | **<0.001** |
| Hypothalamus | Chlorpyrifos concentration | 1.79 | 4 | 0.775 |
|  | Total brain size | 7.05 | 1 | **0.008** |

In the analyses of the fitness-related traits, chronical exposure (i.e. throughout their larval development) to chlorpyrifos had a significant effect on body mass at metamorphosis: individuals exposed to the higher concentration had lower mass than the control individuals (Fig. 3). However, this decrease was not detectable in case of froglets, body mass at dissection was not affected by chlorpyrifos in either concentration (Table 3). Time to metamorphosis (mean ± SE: 40.2 ± 0.69 in the control, 40.1 ± 0.69 in the 0.5 μg/L and 41 ± 0.71 in the 5 μg/L chlorpyrifos treatment) and survival were also not affected by the chlorpyrifos treatments (Table 3). Only seven individuals died during the experiment, six in the higher concentration chronic treatment group (four after finishing the treatment) and one in the control group before metamorphosis.

**Fig. 3.**
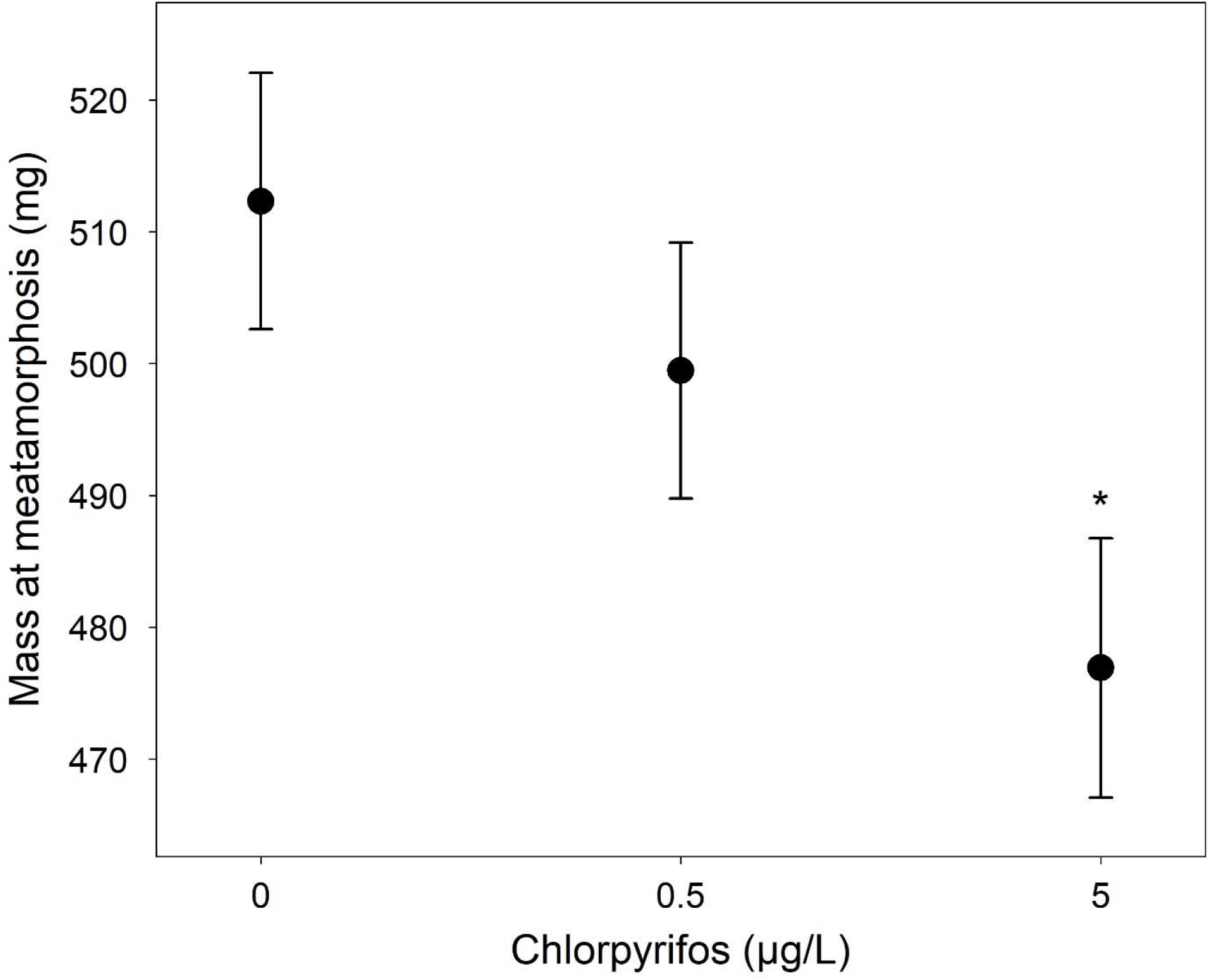
Effects of chlorpyrifos treatments on tadpole mass at metamorphosis. Error bars show the means and standard errors estimated from the model in Table 2.

**Table 3.** Effects of chronic exposure to chlorpyrifos at a concentration of 0.5 and 5 μg/L on the fitness-related traits. Coefficients (*c*) represent the mean in the control group and the differences of the mean in each remaining treatment group from the control group; significant treatment effects (*P* < 0.05) are highlighted in bold.

| Dependent variable | Parameter | *c* | SE | *z* | *P* |
| --- | --- | --- | --- | --- | --- |
| Tadpole survival<sup>1</sup> | Control | -3.46 | 0.84 | -4.12 |  |
|  | Chlorpyrifos 0.5 µg/L | -1.09 | 1.66 | -0.66 | 0.509 |
|  | Chlorpyrifos 5 µg/L | 1.43 | 0.95 | 1.50 | 0.133 |
| Time to metamorphosis<sup>2</sup> | Control | 3.69 | 0.02 | 212.68 |  |
|  | Chlorpyrifos 0.5 µg/L | -0.003 | 0.02 | -0.14 | 0.886 |
|  | Chlorpyrifos 5 µg/L | 0.02 | 0.02 | 0.99 | 0.320 |
| Mass at metamorphosis (mg) | Control | 512.33 | 9.69 | 52.89* |  |
|  | Chlorpyrifos 0.5 µg/L | -12.85 | 13.23 | -0.97* | 0.333 |
|  | Chlorpyrifos 5 µg/L | -35.39 | 13.29 | -2.66* | **0.009** |
| Mass at dissection (g) | Control | -1.83 | 0.18 | -10.31* |  |
|  | Chlorpyrifos 0.5 µg/L | 0.01 | 0.03 | 0.43* | 0.669 |
|  | Chlorpyrifos 5 µg/L | -0.01 | 0.03 | -0.17* | 0.863 |
|  | Days to dissection | 0.03 | 0.002 | 19.54* | **<0.001** |
| Sex ratio | Control | 0.04 | 0.45 | 0.08 |  |
|  | Chlorpyrifos 0.5 µg/L | 0.21 | 0.45 | 0.46 | 0.647 |
|  | Chlorpyrifos 5 µg/L | -0.03 | 0.47 | -0.05 | 0.958 |

<sup>1</sup> logit scale
<sup>2</sup> log scale
\* t-value

We found that sex ratio was independent from the chemical treatments (Table 3). From the 135 individuals that reached the age for phenotypic sexing, we found 23 females and 24 males in the control treatment, 21 females and 26 males in the lower, and 20 females and 21 males in the higher concentration treatment.

## Discussion

In this study, we investigated the effects of larval exposure to ecologically relevant concentrations of chlorpyrifos on the anti-predator behavior, brain morphology, survival, growth and somatic development, and sex ratio of the agile frog. We found that chlorpyrifos had no significant effects on the measured variables, except that chronic exposure to a concentration of 5 μg/L resulted in increased tadpole activity and decreased body mass at metamorphosis. Neither the chronically applied 0.5 μg/L concentration nor the acute 96-h exposure to either of the two concentrations had a significant effect on the traits that we studied.

Tadpole behavior changed upon chronic exposure to chlorpyrifos at a concentration of 5 μg/L: agile frog larvae swam longer distances than their conspecifics raised in pure RSW. Similarly, Richards and Kendall (2003) found that premetamorphic *Xenopus laevis* larvae increased their swimming ability if they were exposed to 1 μg/L or 10 μg/L chlorpyrifos. Another study (Woodley et al., 2015) found that leopard frog (*Lithobates pipiens*) tadpoles tend to increase their activity at 5 μg/L chlorpyrifos concentration, but this effect was not statistically significant. Such intensifications of locomotor activity are probably caused by the moderate AChE inhibition and subsequent accumulation of acetylcholine (Richards and Kendall, 2003; Woodley et al., 2015). Higher activity in larval amphibians may increase predation risk (Skelly, 1994). However, we found no evidence that chlorpyrifos would influence the behavioral response to predator cues in agile frog tadpoles. Surprisingly, the presence of predator chemical cues had only minor effects on the behavior of tadpoles. This result was unexpected because tadpoles usually respond almost instantly to the appearance of chemical cues of predation threat by altering their behavior (Orizaola et al., 2012; Van Buskirk et al., 2014), and the cue concentrations that we applied are known to be perceived by tadpoles and to elicit clear antipredator responses (Hanlon and Relyea, 2013; Winkler and Van Buskirk, 2012), including agile frog larvae (Hettyey et al. 2016). However, such responses are weaker in populations that do not coexist with fishes (Hettyey et al. 2016), which might explain the low responsiveness in our study since the tadpoles used in the study originated from an isolated hill pond lacking fishes. Nevertheless, tadpoles did seem to detect the chemical cues indicating predation risk because they changed their space use in response to it. Because previous studies reported that the detection of chemical cues of predators may be disrupted by chlorpyrifos in fish prey (Maryoung et al., 2015; Sandahl et al., 2004; Tilton et al., 2011), further investigations are required to elucidate whether and under what circumstances may chlorpyrifos influence the anti-predatory responses of amphibians.

Chlorpyrifos did not affect total brain size or the size of any brain region at either concentration. Woodley et al. (2015) exposed leopard frog tadpoles to 5 ppb (0.005 μg/L) chlorpyrifos, which caused narrower and shorter brains compared to the control group, while McClelland et al. (2018) found that exposure to 1 μg/L chlorpyrifos during larval development resulted in brains that were wider than those in the control individuals. It is not surprising that we did not find an effect of chlorpyrifos on brain morphology in the acute treatment groups where tadpoles were exposed to the chemical only for three days. However, the fact that we did not observe any effect on brain morphology in agile frog tadpoles that were exposed chronically to a concentration of 5 μg chlorpyrifos/L is puzzling. Further investigations are needed to understand the causes and consequences of the heterogeneous effects of chlorpyrifos on amphibian neurodevelopment and brain architecture.

We found that chronic exposure to 5 μg/L chlorpyrifos reduced body mass at metamorphosis. Previous studies also found that this chemical can have a negative effect on tadpole mass (Richards and Kendall, 2003; Widder and Bidwell, 2006; Woodley et al., 2015). This suggests that the higher values of ecologically relevant concentrations of chlorpyrifos may decrease population viability, because metamorphic mass is an important determinant of survival to maturity in amphibians (Altwegg and Reyer, 2003; Berven, 1990; Smith, 1987; Üveges et al., 2016). However, the negative effect of chlorpyrifos on metamorphic mass disappeared after ca. two months: body mass at dissection of the chlorpyrifos exposed animals did not differ from the control individuals. This result might be explained with compensatory growth, an accelerated growth which occurs if environmental conditions improve following a period of growth inhibition (Hector et al., 2012; Squires et al., 2010), which often have negative effects later in life due to weaker cellular immune systems and lower total leucocyte numbers (Gervasi and Foufopoulos, 2008).

In our experiment, the lack of significant effects of chlorpyrifos on survival and development were probably due to the use of relatively low concentrations. The majority of previous studies applied much higher concentrations (Barreto et al., 2020; Bernabò et al., 2011a, 2011b; De Arcaute et al., 2012; Richards and Kendall, 2003, 2002; Rutkoski et al., 2020; Silva et al., 2020; Widder and Bidwell, 2008, 2006), although some of these concentrations were referred to as ecologically relevant (Barreto et al., 2020; Bernabò et al., 2011b; Widder and Bidwell, 2008). In these earlier experiments, the applied concentrations were based on surveys from the United States, Canada and South America, where this active ingredient was used in much higher amounts than in Europe e.g. in 2003, 4000-4900 tons of chlorpyrifos were used in the US, while only 1226 tons in European Union (EUROSTAT, 2007; Grube et al., 2006). Studies that used similar concentrations as we did in our experiment also found no significant effects on these variables (Barreto et al., 2020; Bernabò et al., 2011a; Widder and Bidwell, 2008, 2006). This highlights the importance of studying the full range of environmentally relevant concentrations in ecotoxicology.

Chlorpyrifos exposure did not affect sex ratio in our study. Bernabò et al. (2011a) found similar results, although they reported that higher chlorpyrifos concentrations (25-50 μg/L) resulted in intersex condition (testicular oocytes) in some agile frogs. We did not comprehensively screen the testes for oocytes in the present study, but we investigated the gonad histology of five randomly chosen males per chemical treatment, and we found only two intersex animals: one in the control group and one in the 0.5 μg/L chlorpyrifos group. Thus, it seems that the ecologically relevant concentrations that we used are not harmful for sexual development in agile frogs. A low incidence of intersexuality may be a natural phenomenon in young post-metamorphic amphibians (Orton and Tyler, 2015), and is in accordance with the rare occurrence of spontaneous sex reversal we found in agile frogs (Mikó et al., 2021; Nemesházi et al., 2020).

## Conclusions

Taken together, our study revealed that environmentally relevant concentrations of chlorpyrifos are not toxic to agile frog tadpoles in terms of direct mortality, time to metamorphosis, neural and gonadal development. Thus, they appear to be less sensitive to chlorpyrifos and can only be harm under natural conditions in extreme cases, with repeated high-dose applications. However, larvae of different amphibian species may largely vary in their susceptibility to pesticides (Bókony et al., 2020; Bridges and Semlitsch, 2000), so it is important to note that ecotoxicological effects cannot be generalized even across relatively closely related species. Furthermore, our results highlight the importance of ecotoxicological studies using concentrations that are based on environmental surveys, because only these can provide realistic assessments of ecological risk.

## Supporting information

Electronic supplementary material

## Acknowledgements

We thank all members of the Lendület Evolutionary Ecology Research Group for insightful discussions, Viktória Verebélyi, Patrik Katona, László Sipőcz and Mátyás Szin for help with animal handling and data archiving, Gergely Zachar for the fixative solution, and Gergő Tholt and the NÖVI Department of Zoology for allowing us to use their stereomicroscope and camera and for providing helpful advice. We thank Bálint Bombay for the paintings of agile frogs.

## Funding

The study was funded by the National Research, Development and Innovation Office of Hungary (NKFIH 115402) and the ‘Lendület’ programme of the Hungarian Academy of Sciences (MTA, MTA, LP2012□24/2012). VB and AH were supported by the János Bolyai Scholarship of the MTA, VV and EN by the Ministry of Human Capacities (National Program for Talent of Hungary, NTP-NFTÖ-18-B-0412 to VV, NTP-NFTÖ-17-B-0317 to EN), and NU by the Young Researcher program of the Hungarian Academy of Sciences. None of the funding sources had any influence on the study design, collection, analysis, and interpretation of data, writing of the paper, or decision to submit it for publication.

## Author contributions

**Zsanett Mikó:** Conceptualization, Methodology, Investigation, Data curation, Formal analysis, Writing - original draft, Writing - review & editing, Visualization. **Veronika Bókony:** Investigation, Formal analysis, Writing - review & editing, Supervision, Funding acquisition. **Nikolett Ujhegyi:** Investigation, Data curation, Writing - review & editing. **Edina Nemesházi:** Investigation, Writing - review & editing. **Réka Erös:** Methodology, Investigation, Data curation, Writing - review & editing. **Stephanie Orf:** Data curation, Writing - review & editing. **Attila Hettyey:** Conceptualization, Methodology, Investigation, Writing - review & editing, Supervision, Funding acquisition.

## Notes

### Competing Interest Statement

The authors have declared no competing interest.

