## Supplementary material for "Effects of chlorpyrifos on early development and anti-predator behavior of agile frogs": Electronic supplementary material

**Table S1.** Parameter estimates (b, with SE) of the general linear mixed effects models for each behavioral variable. Intercept refers to the control group (no chlorpyrifos, no predator cue), first observation day. Baseline behavior means the first 20 minutes of recording without predator cue. Time of day is the observation session, ranging from 1 to 7.

| Behavioral variable | Fixed effects | b | SE | df | *t* | *P* |
| --- | --- | --- | --- | --- | --- | --- |
| Swimming trajectory length | Intercept | 4.392 | 1.826 | 278.269 | 2.405 | 0.017 |
|  | Chlorpyrifos 0.5 μg/L acute | -2.422 | 2.047 | 318.272 | -1.183 | 0.238 |
|  | Chlorpyrifos 5 μg/L acute | -0.619 | 2.015 | 311.921 | -0.308 | 0.759 |
|  | Chlorpyrifos 0.5 μg/L chronic | 0.042 | 2.161 | 307.794 | 0.019 | 0.985 |
|  | Chlorpyrifos 5 μg/L chronic | 3.183 | 2.013 | 314.220 | 1.581 | 0.115 |
|  | Day 2 | -1.867 | 1.692 | 167.364 | -1.104 | 0.271 |
|  | Predator cue | -1.918 | 2.110 | 306.918 | -0.909 | 0.364 |
|  | Baseline behavior | 0.405 | 0.032 | 348.036 | 12.838 | **<0.001** |
|  | Time of day | 0.481 | 0.193 | 150.645 | 2.491 | 0.014 |
|  | Chlorpyrifos 0.5 μg/L acute × Day 2 | 2.230 | 2.417 | 172.671 | 0.923 | 0.358 |
|  | Chlorpyrifos 5 μg/L acute × Day 2 | -0.922 | 2.377 | 168.162 | -0.388 | 0.699 |
|  | Chlorpyrifos 0.5 μg/L chronic × Day 2 | 0.736 | 2.532 | 166.429 | 0.291 | 0.772 |
|  | Chlorpyrifos 5 μg/L chronic × Day 2 | -0.942 | 2.368 | 170.026 | -0.398 | 0.691 |
|  | Chlorpyrifos 0.5 μg/L acute × Predator cue | 3.346 | 2.997 | 323.422 | 1.116 | 0.265 |
|  | Chlorpyrifos 5 μg/L acute × Predator cue | 4.588 | 2.903 | 314.655 | 1.580 | 0.115 |
|  | Chlorpyrifos 0.5 μg/L chronic × Predator cue | 2.003 | 2.951 | 318.232 | 0.679 | 0.498 |
|  | Chlorpyrifos 5 μg/L chronic × Predator cue | -0.351 | 2.957 | 318.224 | -0.119 | 0.906 |
|  | Day 2 × Predator cue | 3.173 | 2.429 | 170.921 | 1.306 | 0.193 |
|  | Chlorpyrifos 0.5 μg/L acute × Day 2 × Predator cue | -4.623 | 3.511 | 174.444 | -1.317 | 0.189 |
|  | Chlorpyrifos 5 μg/L acute × Day 2 × Predator cue | -3.358 | 3.442 | 171.189 | -0.976 | 0.331 |
|  | Chlorpyrifos 0.5 μg/L chronic × Day 2 × Predator cue | -2.877 | 3.441 | 168.504 | -0.836 | 0.404 |
|  | Chlorpyrifos 5 μg/L chronic × Day 2 × Predator cue | 0.739 | 3.465 | 170.078 | 0.213 | 0.831 |
| Proportion of time near wall | Intercept | 52.961 | 6.512 | 359.777 | 8.132 | **<0.001** |
|  | Chlorpyrifos 0.5 μg/L acute | 4.529 | 6.165 | 361.539 | 0.735 | 0.463 |
|  | Chlorpyrifos 5 μg/L acute | -6.871 | 6.089 | 361.487 | -1.128 | 0.260 |
|  | Chlorpyrifos 0.5 μg/L chronic | 2.967 | 6.534 | 361.489 | 0.454 | 0.650 |
|  | Chlorpyrifos 5 μg/L chronic | 2.361 | 6.029 | 361.329 | 0.392 | 0.696 |
|  | Day 2 | -5.289 | 5.966 | 175.944 | -0.886 | 0.377 |
|  | Predator cue | -0.669 | 6.251 | 361.493 | -0.107 | 0.915 |
|  | Baseline behavior | 0.327 | 0.059 | 358.671 | 5.482 | **<0.001** |
|  | Time of day | -0.255 | 0.535 | 180.910 | -0.477 | 0.634 |
|  | Chlorpyrifos 0.5 μg/L acute × Day 2 | 3.953 | 8.511 | 183.589 | 0.464 | 0.643 |
|  | Chlorpyrifos 5 μg/L acute × Day 2 | 6.575 | 8.395 | 178.313 | 0.783 | 0.434 |
|  | Chlorpyrifos 0.5 μg/L chronic × Day 2 | 1.775 | 8.954 | 176.114 | 0.198 | 0.843 |
|  | Chlorpyrifos 5 μg/L chronic × Day 2 | -0.879 | 8.348 | 180.142 | -0.105 | 0.916 |
|  | Chlorpyrifos 0.5 μg/L acute × Predator cue | -10.693 | 9.049 | 361.620 | -1.182 | 0.238 |
|  | Chlorpyrifos 5 μg/L acute × Predator cue | -2.778 | 8.782 | 361.546 | -0.316 | 0.752 |
|  | Chlorpyrifos 0.5 μg/L chronic × Predator cue | -3.769 | 8.861 | 361.543 | -0.425 | 0.671 |
|  | Chlorpyrifos 5 μg/L chronic × Predator cue | -9.489 | 8.888 | 361.294 | -1.068 | 0.286 |
|  | Day 2 × Predator cue | 2.554 | 8.559 | 181.049 | 0.298 | 0.766 |
|  | Chlorpyrifos 0.5 μg/L acute × Day 2 × Predator cue | -0.086 | 12.348 | 185.630 | -0.007 | 0.994 |
|  | Chlorpyrifos 5 μg/L acute × Day 2 × Predator cue | 3.402 | 12.139 | 181.649 | 0.280 | 0.780 |
|  | Chlorpyrifos 0.5 μg/L chronic × Day 2 × Predator cue | -1.018 | 12.139 | 178.429 | -0.084 | 0.933 |
|  | Chlorpyrifos 5 μg/L chronic × Day 2 × Predator cue | 0.281 | 12.221 | 180.403 | 0.023 | 0.982 |
| Mobility rate of the tadpole | Intercept | 30.614 | 3.862 | 337.491 | 7.927 | **<0.001** |
|  | Chlorpyrifos 0.5 μg/L acute | -3.177 | 1.921 | 334.349 | -1.653 | 0.099 |
|  | Chlorpyrifos 5 μg/L acute | -0.506 | 1.891 | 330.182 | -0.267 | 0.789 |
|  | Chlorpyrifos 0.5 μg/L chronic | 0.333 | 2.016 | 324.701 | 0.165 | 0.869 |
|  | Chlorpyrifos 5 μg/L chronic | 0.319 | 1.897 | 336.487 | 0.168 | 0.866 |
|  | Day 2 | 0.090 | 1.678 | 179.042 | 0.054 | 0.957 |
|  | Predator cue | -1.981 | 2.007 | 337.807 | -0.987 | 0.324 |
|  | Baseline behavior | 0.532 | 0.047 | 345.910 | 11.300 | **<0.001** |
|  | Time of day | 0.083 | 0.173 | 161.946 | 0.480 | 0.632 |
|  | Chlorpyrifos 0.5 μg/L acute × Day 2 | 2.257 | 2.402 | 186.432 | 0.940 | 0.349 |
|  | Chlorpyrifos 5 μg/L acute × Day 2 | -0.602 | 2.362 | 181.062 | -0.255 | 0.799 |
|  | Chlorpyrifos 0.5 μg/L chronic × Day 2 | -1.338 | 2.517 | 179.052 | -0.532 | 0.596 |
|  | Chlorpyrifos 5 μg/L chronic × Day 2 | 0.585 | 2.355 | 183.408 | 0.248 | 0.804 |
|  | Chlorpyrifos 0.5 μg/L acute × Predator cue | 4.495 | 2.806 | 332.487 | 1.602 | 0.110 |
|  | Chlorpyrifos 5 μg/L acute × Predator cue | 2.776 | 2.720 | 328.488 | 1.020 | 0.308 |
|  | Chlorpyrifos 0.5 μg/L chronic × Predator cue | 1.931 | 2.753 | 330.859 | 0.701 | 0.484 |
|  | Chlorpyrifos 5 μg/L chronic × Predator cue | 4.361 | 2.771 | 332.442 | 1.573 | 0.117 |
|  | Day 2 × Predator cue | 1.239 | 2.416 | 184.352 | 0.513 | 0.609 |
|  | Chlorpyrifos 0.5 μg/L acute × Day 2 × Predator cue | -1.977 | 3.493 | 188.957 | -0.566 | 0.572 |
|  | Chlorpyrifos 5 μg/L acute × Day 2 × Predator cue | -0.175 | 3.417 | 184.179 | -0.051 | 0.959 |
|  | Chlorpyrifos 0.5 μg/L chronic × Day 2 × Predator cue | 0.854 | 3.418 | 181.357 | 0.250 | 0.803 |
|  | Chlorpyrifos 5 μg/L chronic × Day 2 × Predator cue | -1.771 | 3.453 | 184.373 | -0.513 | 0.609 |
